# Context-dependent extinction learning emerging from raw sensory inputs: A reinforcement learning approach

**DOI:** 10.1101/2020.04.27.059121

**Authors:** Thomas Walther, Nicolas Diekmann, Sandhiya Vijayabaskaran, José R. Donoso, Denise Manahan-Vaughan, Laurenz Wiskott, Sen Cheng

**Affiliations:** Institute for Neural Computation, Ruhr University Bochum, Germany; Neurophysiology, Medical Faculty, Ruhr University Bochum, Germany

## Abstract

The context-dependence of extinction learning has been well studied and requires the hippocampus. However, the underlying neural mechanisms are still poorly understood. Using memory-driven reinforcement learning and deep neural networks, we developed a model that learns to navigate autonomously in biologically realistic VR environments based on raw camera inputs alone. Neither is context represented explicitly in our model, nor is context change signaled. We find that memory-intact agents learn distinct context representations, and develop ABA renewal, whereas memory-impaired agents do not. These findings reproduce the behavior of control and hippocampal animals, respectively. We therefore propose that the role of the hippocampus in the context-dependence of extinction learning might stem from its function in episodic-like memory and not in context-representation per se. We conclude that context-dependence can emerge from raw visual inputs.

## Introduction

Treatment of anxiety disorders by exposure therapy is often followed by unwanted renewal of the seemingly extinguished fear. A better understanding of the cognitive and neural mechanisms governing extinction learning and fear renewal is therefore needed to develop novel therapies. The study of extinction learning goes back to Ivan Pavlov’s research on classical conditioning [1]. Pavlov and his colleagues discovered that the conditioned response (CR) to a neutral conditioned stimulus (CS) diminishes over time, if the CS is presented repeatedly without reinforcement by an unconditioned stimulus (US). This phenomenon is called extinction learning.

The arguably best understood extinction learning paradigm is fear extinction, in which an aversive stimulus, e.g. an electric foot shock, is used as US [2]. The conditioned fear response diminishes during extinction, only to return later under certain circumstances [3]. This effect is known as fear renewal, and can be controlled by context changes, as shown in aversive ABA renewal: Subjects acquire a fear response to a CS, e.g. a neutral tone, by repeatedly coupling it with an US, e.g. a foot shock, in context A. Then the fear response is extinguished in a different context B, and eventually tested again in context A in the absence of the US. Renewal means that the fear response returns in the test phase despite the absence of the US [4]. ABA renewal demonstrates that extinction learning does not delete the previously learned association, although it might weaken the association [3]. Furthermore, ABA renewal underscores that extinction learning is strongly context-dependent [4–10].

Fear extinction is thought to depend mainly on three cerebral regions: The amygdala, responsible for initial fear learning, the ventral medial prefrontal cortex, guiding the extinction process, and the hippocampus, controlling the context-specific retrieval of extinction [3] and/or encoding contextual information [11]. Here, we focus on the role of the hippocampus in extinction learning tasks. Lesions or pharmacological manipulations of the hippocampus have been reported to induce deficits in context encoding [3, 11], which in turn impede context disambiguation during extinction learning [5–8]. In particular, André and Manahan-Vaughan (2016) found that intracerebral administration of a dopamine receptor agonist, which alters synaptic plasticity in the hippocampus [12], reduced the context-dependent expression of renewal in an appetitive ABA renewal task in a T-maze [9]. This task depends on the the specific involvement of certain subfields of the hippocampus [13].

However, it remains unclear how behavior becomes context-dependent and what neural mechanisms underlie this learning. The majority of computational models targets exclusively extinction tasks that involve classical conditioning, like the earliest model of conditioning, the Rescorla-Wagner model [14] (for an overview, see [3]). However, in its original form this model treats extinction learning as unlearning of previously acquired associations between CS and US and so could not account for the phenomenology of extinction learning, including renewal in particular. Nevertheless, the Rescorla-Wagner model inspired the development of reinforcement learning (RL) and many recent models of extinction learning. For instance, Ludvig et al (2017) added experience replay [15] to the Rescorla-Wagner model to account for a range of phenomena associated with extinction in classical conditioning: spontaneous recovery, latent inhibition, retrospective revaluation, and trial spacing effects [16]. However, their approach required explicit signals for cues and context, and did not learn from realistic visual information. Moreover, Ludvig et al. acknowledged that their method required the integration of more advanced RL techniques in order to handle operant conditioning tasks.

Computational models of extinction learning based on operant/instrumental conditioning are rather rare [3]. One of the few models was developed by Redish et al. and was based on RL driven by temporal differences. Their model emphasized the importance of a basic memory system to successfully model renewal in operant extinction tasks [17]. However, their model relied on the existence of dedicated signals for representing cues and contexts, and did not learn from realistic visual input.

Overcoming the caveats of existing approaches, we propose a novel model of operant extinction learning that combines deep neural networks with experience replay memory and reinforcement learning, the deep Q-network (DQN) approach [15]. Our model does not require its architecture to be manually tailored to each new scenario and does not rely on the existence of explicit signals representing cues and contexts. It can learn complex spatial navigation tasks based on raw visual information, and can successfully account for instrumental ABA renewal without receiving a context signal from external sources. Our model also shows better performance than the model by Redish et al.: faster acquisition in simple mazes, and more biologically plausible extinction learning in a context different from the acquisition context. In addition, the deep neural network in our model allows for studying the internal representations that are learned by the model. Distinct representations for contexts A and B emerge in our model during extinction learning from raw visual inputs obtained in biologically plausible virtual reality (VR) scenarios and reward contingencies.

## Materials and Methods

### Modeling Framework

All simulations in this paper were performed in a VR testbed designed to study biomimetic models of rodent behavior in spatial navigation and operant extinction learning tasks (Fig. 1). The framework consists of multiple modules, including a VR component based on the Blender rendering and simulation engine (The Blender Foundation, https://www.blender.org/), in which we built moderately complex and biologically plausible virtual environments. In the VR, the RL agent is represented by a small robot that is equipped with a panoramic camera to simulate the wide field of view of rodents.

**Fig 1.**
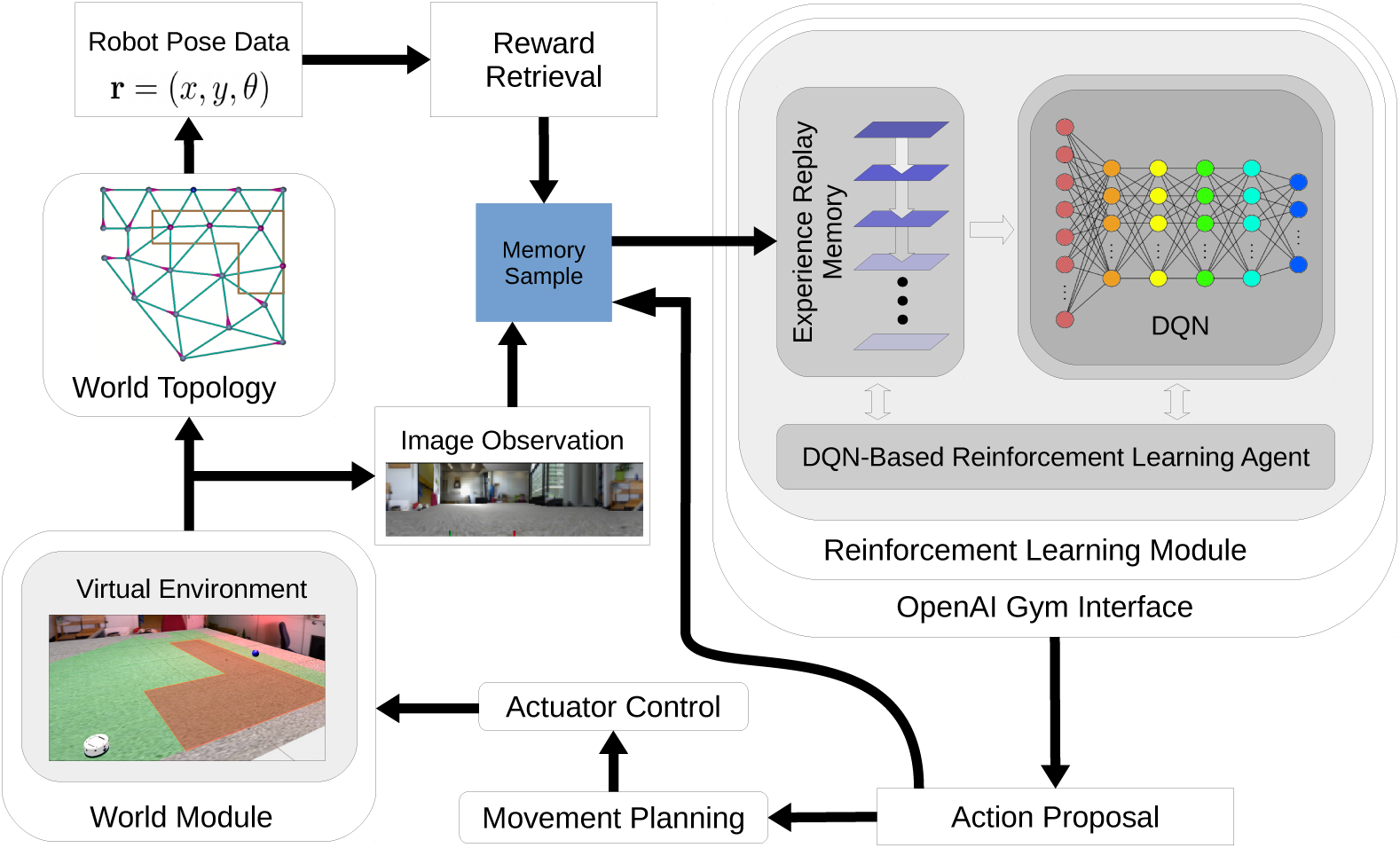
Model overview. Modules are sketched as rounded rectangles, other rectangles represent data and commands, the information/control flow is indicated by arrows. See the Materials and Methods for an explanation of our model.

The modeling framework allows for two modes of operation: an online model, in which simulations are run in real-time and physics-based simulations are supported, and an offline mode, in which multiple simulations can be run in parallel on pre-rendered camera images. The latter operating mode increases the computational efficiency of our model, and facilitates the rapid generation of large numbers of learning trials and repetitions. All results in this paper were obtained using the offline mode.

### Visually Driven Reinforcement Learning with Deep Q-Networks

The agent learns autonomously to navigate in an environment using RL [18]: Time is discretized in time steps *t*; the state of the environment is represented by 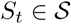, where 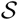 is the set of all possible states. The agent observes the state *S*_*t*_ and selects an appropriate action,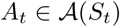, where 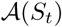 is the set of all possible actions in the current state. In the next time step, the environment evolves to a new state *S*_*t*+1_, and the agent receives a reward *R*_*t*+1_.

In our modeling framework, the states correspond to spatial positions of the agent within the VR. The agent observes these states through visual images captured by the camera from the agent’s position (e.g. Fig. 2). The set of possible states, 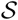, is defined by the discrete nodes in a manually constructed topology graph that spans the virtual environment (Fig. 3). State space discretization accelerates the convergence of RL. The possible actions in each state, 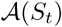, are defined by the set of outgoing connections from the node in the topology graph. Once an action is chosen, the agent moves to the connected node. In the current implementation, the agent does not rotate.

**Fig 2.**
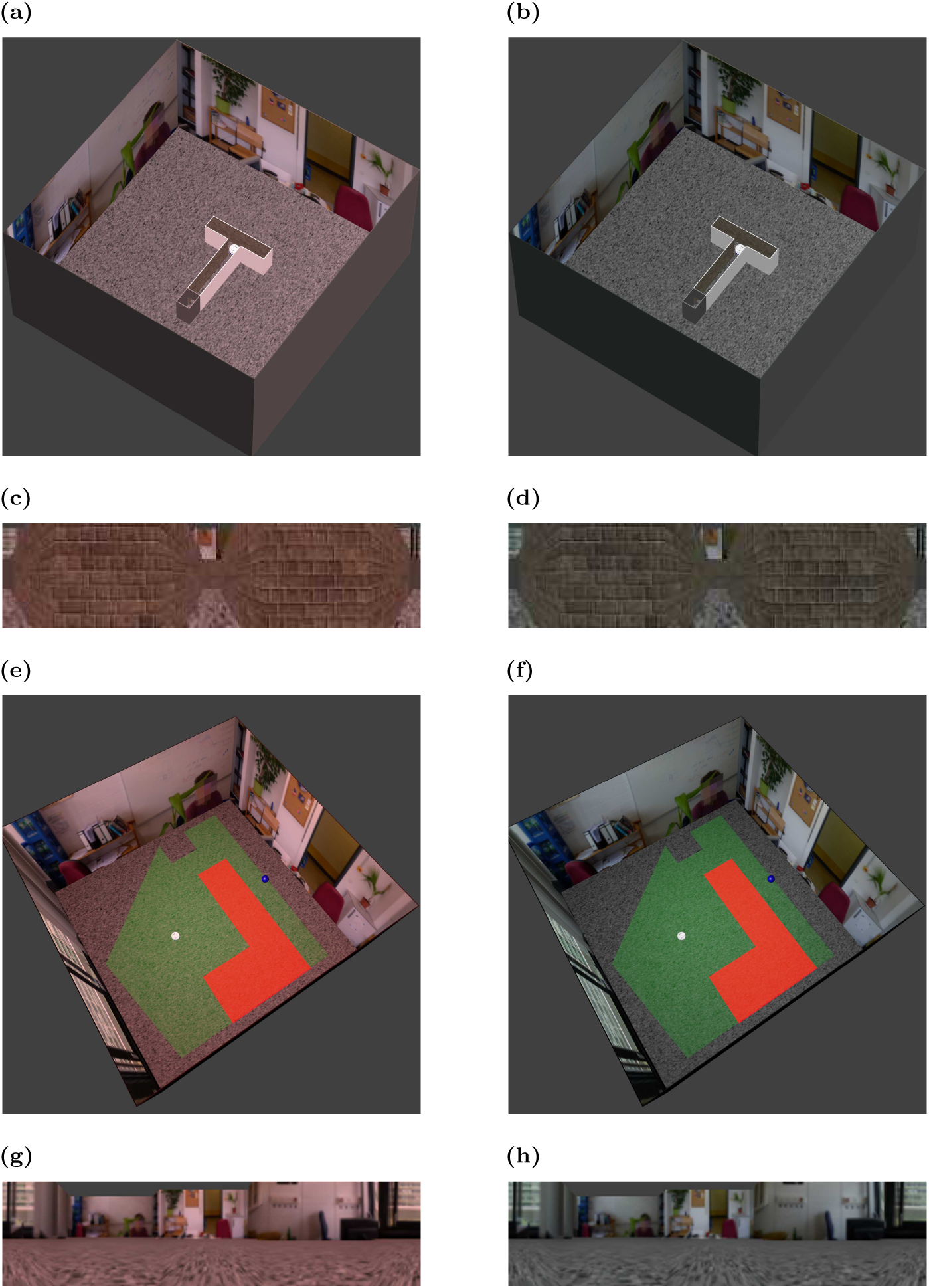
Examples of virtual environments and camera views. **(a,b)** Overview of T-maze in two different contexts. Two different contexts are generated by using different colors for the overall illumination. **(c,d)** Camera views in T-maze from the agent’s perspective. **(e,f)** Overview of shock zone maze used to study inhibitory avoidance. **(g,h)** Camera views in the shock zone maze.

**Fig 3.**
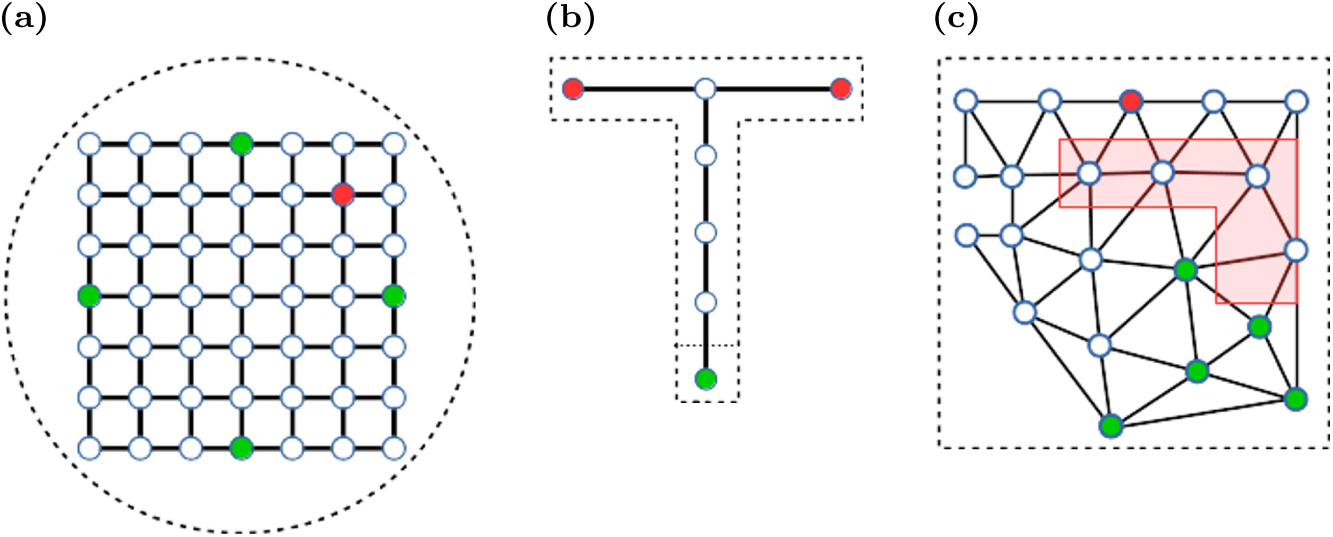
Topology graphs for different simulation environments. **(a)** Morris watermaze. **(b)** T-maze. **(c)** Shock zone maze. The red polygon indicates the shock zone. Nodes in the graph indicate allowed positions, and solid lines allowed transitions between positions. Start nodes are marked green, and (potential) goal nodes are marked red. The dashed lines sketch the maze perimeter.

For the learning algorithm we adopt a specific variant of RL called Q-learning, where the agent learns a state-action value function *Q* (*S, A*), which represents the value to the agent of performing action *A* in state *S*. The agent learns from interactions with the environment by updating the state-action value function according to the following rule [18]:

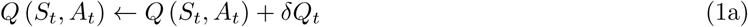

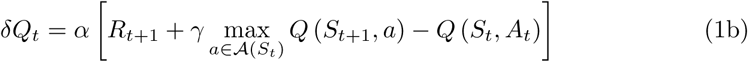

where *α* is the learning rate and *γ* is the discount factor that devalues future rewards relative to immediate ones. The Q-learning update rule has been shown mathematically to converge to a local optimum. Since the state-action value function is likely very complex and its analytical model class is unknown, we use a deep neural network to approximate *Q*(*S, A*) [15]. In our simulations, the DQN consists of fully connected, dense layers between the input and output layer and the weights are initialized with random values. We use the tanh-function as the activation function.

The input to the network consists of (30,1)-pixel RGB images, which are captured in the virtual environment. The network architecture comprises an input layer (with 90 neurons for 30 pixels x 3 channels), four hidden layers (64 neurons each), that enable the DQN to learn complex state-action value functions, and an output layer that contains a single node for each action the agent can perform. We use the Keras^1^ and KerasRL^2^ implementation for the DQN in our model.

To balance between exploration and exploitation our agent uses an *ϵ*-greedy strategy [18]: In each state *S*_*t*_, the agent either selects the action that yields the highest *Q* (*S*_*t*_, *A*_*t*_) with a probability of 1 − *ϵ* (exploitation), or chooses a random element from the set of available actions 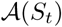 with a probability of *ϵ* (exploration). To ensure sufficient exploration, we set *ϵ* = 0.3 in all simulations, unless otherwise noted.

In our model, experiences are stored in memory to be replayed later. An experience contains information about the state observed, action taken, and rewards received, i.e. (*S*_*t*_, *A*_*t*_, *S*_*t*+1_, *R*_*t*+1_). The memory module uses first-in-first-out memory, i.e. if the memory’s capacity *c* is exceeded, the oldest experiences are deleted first to make room for novel ones. In each time step, random batches of memory entries are sampled for experience replay [15]. This experience replay in RL was inspired by hippocampal replay of sequential neural activity encountered during experience, which is hypothesized to play a critical role in memory consolidation and learning [19, 20]. We therefore suggest that lesions or drug-induced impairments of the hippocampal formation can be simulated by reducing the agent’s replay memory capacity.

Positive or negative rewards drive RL [18]. These rewards are task-specific and will be defined separately for each simulation setup below.

### Operant Learning Tasks Simulated in our Model

#### General Setup

All trials used in our simulations had a common structure. Each trial began with the agent located at one of the defined starting nodes. In each following time step, the agent could chose to execute one action that was allowed in the current state as defined by the topology graph. Each time step incurred a negative reward of −1 to encourage the agent to find the shortest path possible from the start to the goal node. Trials ended when either the agent reached the goal node, or the number of simulated time steps exceeded 100 (time-out). The experience replay memory capacity was set to *c* = 5000. Different contexts were defined by the illumination color of the house lights. Context A was illuminated with red house lights and context B with white house lights.

#### Linear Track

As a proof-of-concept and to facilitate a comparison to Redish et al.’s model, we first studied spatial learning on a simple linear track. The track was represented by a one-dimensional topology graph consisting of *N* = 3,…, 15 nodes. Since Redish et al.’s model requires clearly separable state representations, we used binary encoded node state indices in this simulation, instead of the raw camera input as in the other simulations. The starting node was always the first node, while the target node was node *N*. The agent received a reward of +1.0 for reaching the target node.

#### Morris watermaze

Next, we simulated spatial navigation in the more challenging Morris watermaze [21] to show that our model can learn operant spatial tasks in 2-d from raw visual input. The watermaze was circular with a diameter of 2 *m*. It was placed in the center of a typical lab environment to provide distal cues. The topology graph for this maze consisted of a rectangular grid that covered the center part of the watermaze (Fig. 3a). Four nodes {N(orth),E(ast),S(outh),W(est)} were used as potential starting nodes. They were selected in analogy to a protocol introduced by Anisman et al. [22]: Trials were arranged into blocks of 4 trials. Within each block, the sequence of starting locations was a random permutation of the sequence (N,E,S,W). The goal node was always centered in the NE quadrant and yielded an additional reward of +1.0.

#### Extinction Learning in T-maze

To study the emergence of context-dependent behavior, we simulated the following ABA renewal experiment [9]: In a simple T-Maze, rats were rewarded for running to the end of one of the side arms (target arm). The acquisition context A was defined by a combination of distal and proximal visual cues, as well as odors. Once the animals performed to criterion, the context was changed to B, which was distinct from A, and the extinction phase began. In this phase, animals were no longer rewarded for entering the former target arm. Once the animals reverted to randomly choosing between the two side arms, the context was switched back to A and the test phase started. Control animals showed renewal and chose the former target arm more frequently than the other arm, even though it was no longer rewarded.

In our modeling framework, we maintained the spatial layout of the experimental setup and the richness of visual inputs in a laboratory setting, which is usually oversimplified in other computational models. A skybox showed a standard lab environment with distal cues for global orientation, and maze walls were textured using a standard brick texture (figs. 2a-2d). We made a few minor adjustments to accommodate the physical differences between rats and the simulated robot. For instance, we had to adjust the size of the T-maze and use different illumination to distinguish contexts A and B. In the rat experiments, the two contexts were demarcated by odor cues in addition to visual cues [9]. If anything, the simplifications in our model make the distinction between contexts harder, since odor cues provide animals a separate channel of information that is irrelevant to the navigation task.

The topology graph in the T-Maze consisted of seven nodes (Fig. 3b). Reaching a node at the end of the side arms yielded a reward of +1.0 to encourage goal-directed behavior in the agent. The left endpoint was the target node and yielded an additional reward of +10.0 only in the acquisition phase, not in the extinction and test phases. As in the experiments, a trial was scored as correct, if the agent entered the target arm, and as incorrect, if the other arm was selected or the trial timed out. Finally, to compare our model account of ABA renewal to that of Redish et al. [17], we used the same topology graph, but provided their model with inputs that represented states and contexts as binary encoded indices.

A simulated session consisted of 300 trials in each of the experimental phases: acquisition, extinction and test. So, altogether a session consisted of 900 trials.

#### Extinction Learning in Inhibitory Avoidance

To model learning in a task that requires complex spatial behavior in addition to dynamically changing reinforcements, we use an inhibitory avoidance task in an ABA renewal design. The agent had to navigate from a variable starting location to a fixed goal location in the shock zone maze (Figs. 2e-h), while avoiding a certain region of the maze, the shock zone. Distal cues were provided in this environment using the same skybox as in the watermaze simulation. The topology graph was built using Delaunay triangulation (Fig. 3c). The start node was randomly chosen from four nodes (green dots). Reaching the goal node (red dot) earned the agent a reward of +1.0. If the agent landed on a node in the shock zone (red area, Figs. 2e-f,3c), it would receive a punishment (reward= −20.0). In both the extinction and test phases, the shock zone was inactive and no negative reward was added for landing on a node within the shock zone.

### Neural Representations of Context in ABA Renewal Tasks

To study the influence of experience replay and model the effect of hippocampal lesions/inactivations, we compared agents with intact memory (*c* = 5000) to memory-impaired agents with *c* = 150.

To study the representations in the DQN layers, we used dimensionality reduction as follows: We defined *I*_*l*_ (*z*, **x**) to represent the activation of layer *l* ∈ [0, 1,…, 5] when the DQN is fed with an input image captured at location **x** in context *z* ∈ {*A, B*}, where *l* = 0 is the input layer, and *l* = 5 the output layer. Let **X**_*A*_ and **X**_*B*_ be the sets of 1000 randomly sampled image locations in context A and B, respectively. We expected that the representations *I*_*l*_ (*A*, **X**_*A*_) and *I*_*l*_ (*B*, **X**_*B*_) in the DQN’s higher layers form compact, well-separated clusters in activation space by the end of extinction. This would enable agents to disambiguate the contexts and behave differently in the following test phase in context A from how they behaved in context B at the end of extinction, which is exactly renewal. By contrast, for memory-impaired agents we would expect no separate clusters for the representations of the two contexts, which would impair the agent’s ability to distinguish between contexts A and B, which in turn would impair renewal. We applied Linear Discriminant Analysis (LDA) to find a clustering that maximally separates *I*_*l*_ (*A*, **X**_*A*_) and *I*_*l*_ (*B*, **X**_*B*_). The cluster distances *d*_*l*_ are related to the context discrimination ability of network layer *l*.

## Results

We developed a computational framework based on reinforcement learning driven by experience memory replay and a deep neural network to model complex operant learning tasks in realistic environments. Unlike previous computational models, our computational modeling framework learns based on raw visual inputs and does not receive pre-processed inputs.

### Proof-of-principle Studies of Spatial Learning

As a proof-of-principle and to facilitate a direct comparison to an earlier model, we first studied spatial learning on a simple linear track with simplified inputs, i.e. the positions were represented by binary indices. To vary the difficulty of the task, the track was represented by a 1-d topology graph with variable node count. The first node was always the starting node and the agent was rewarded for reaching the last node. The larger the number of nodes in the graph, the more difficult it was for the agent to solve the task. Hence, the average number of trials, in which the agent reached the goal node before the trial timed out, decreased with the number of nodes *N* (Fig. 4a, blue line). The reason for this result is simple. The agent had to find the goal node by chance at least once before it could learn a goal-directed behavior. This was less likely to occur in longer tracks with a larger number of intermediate nodes.

**Fig 4.**
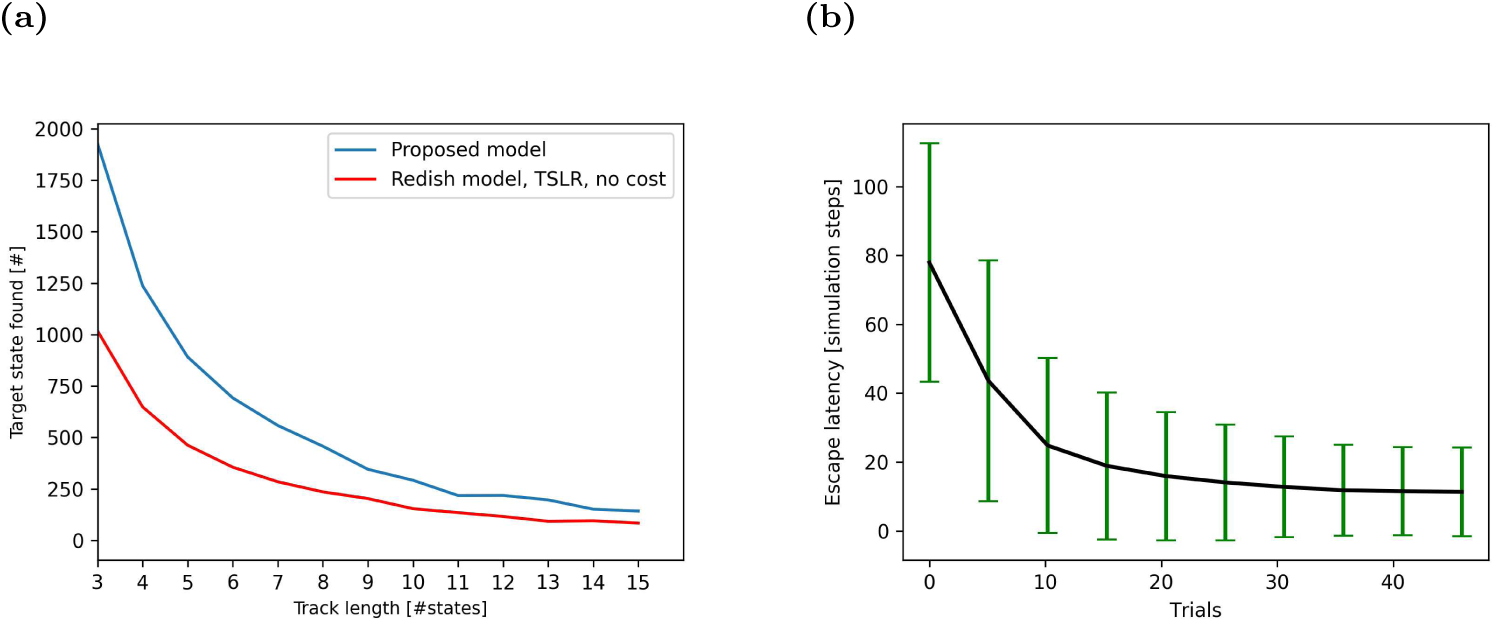
Performance on spatial navigation tasks. **(a)** Number of times the agent finds the goal location given a total of 5000 simulation steps on a linear track in our model (blue line) and comparison model [17] (red line). Results are averaged over 50 simulations for each track length. **(b)** Escape latency in the virtual watermaze decreases with learning trials. Data averaged over 1000 agents, error bars represent standard deviations.

Comparing our model to existing approaches turned out to be difficult, since operant extinction learning has not received much attention in the past. The model proposed by Redish et al. [17] was eventually chosen as a reference. Even though this model was developed for different kinds of tasks, it could solve the spatial learning task on the linear track if inputs provided to the model were simplified to represent states as indices (Fig. 4b, red line). However, Redish et al.’s model did not perform as well as our model regardless of the track length. This result is due to experience memory in our model, which could replay samples of successful navigation to speed up learning.

Next, we studied more complex spatial learning in a 2-d watermaze [21] with raw camera inputs. Since the agent was rewarded for finding the location of the escape platform as quickly as possible, we quantified learning performance by the escape latency (Fig. 4a). The learning curve indicates clearly that the system’s performance improved rapidly with learning trials, i.e. the latency decreased. Our result compared well to those reported in the literature on rodents [23].

### Extinction and Renewal of Operant Behavior

Having shown that our model agent can learn simple spatial navigation behaviors based on raw camera inputs, we turned our attention to more complex learning experiments such as operant extinction and renewal. Initial tests were run in the T-maze. Performance was measured with cumulative response curves (CRCs). If the agent entered the target arm, the CRC was increased by one. If the other arm was entered or the trial timed out, the CRC was not changed. Hence, consistent correct performance yielded a slope of 1, (random) alternation of correct and incorrect responses yielded a slope of 0.5, and consistent incorrect responses yielded a slope of 0. The CRC for our model looked just as expected for an ABA renewal experiment (Fig. 5a). After a very brief initial learning period, the slope increased to about 1 until the end of the acquisition phase, indicating nearly perfect performance. At the transition from the acquisition to the extinction phase, the slope of the CRCs stayed constant in the first 50 extinction trials, indicating that the agent generalized the learned behavior from context A to context B, even though it no longer received a larger reward for the left turn. After 50 extinction trials, the slope decreased gradually (Fig. 5a, red arrow), indicating extinction learning. Finally, the change from context B back to context A at the onset of the test phase triggered an abrupt rise of the CRC slope, indicating a renewal of the initially learned behavior. The CRC for Redish et al.’s model looked similar to that for our model (Fig. 5b), but exhibited two key differences that were not biologically plausible. First, the CRCs split into one group that showed no extinction, and hence no renewal either, and another group that showed extinction and renewal. Second, the latter group showed a sharp drop of the CRC slope in the very first extinction trial (Fig. 5b, red arrow), indicating an immediate shift in behavior upon the first unrewarded trial.

**Fig 5.**
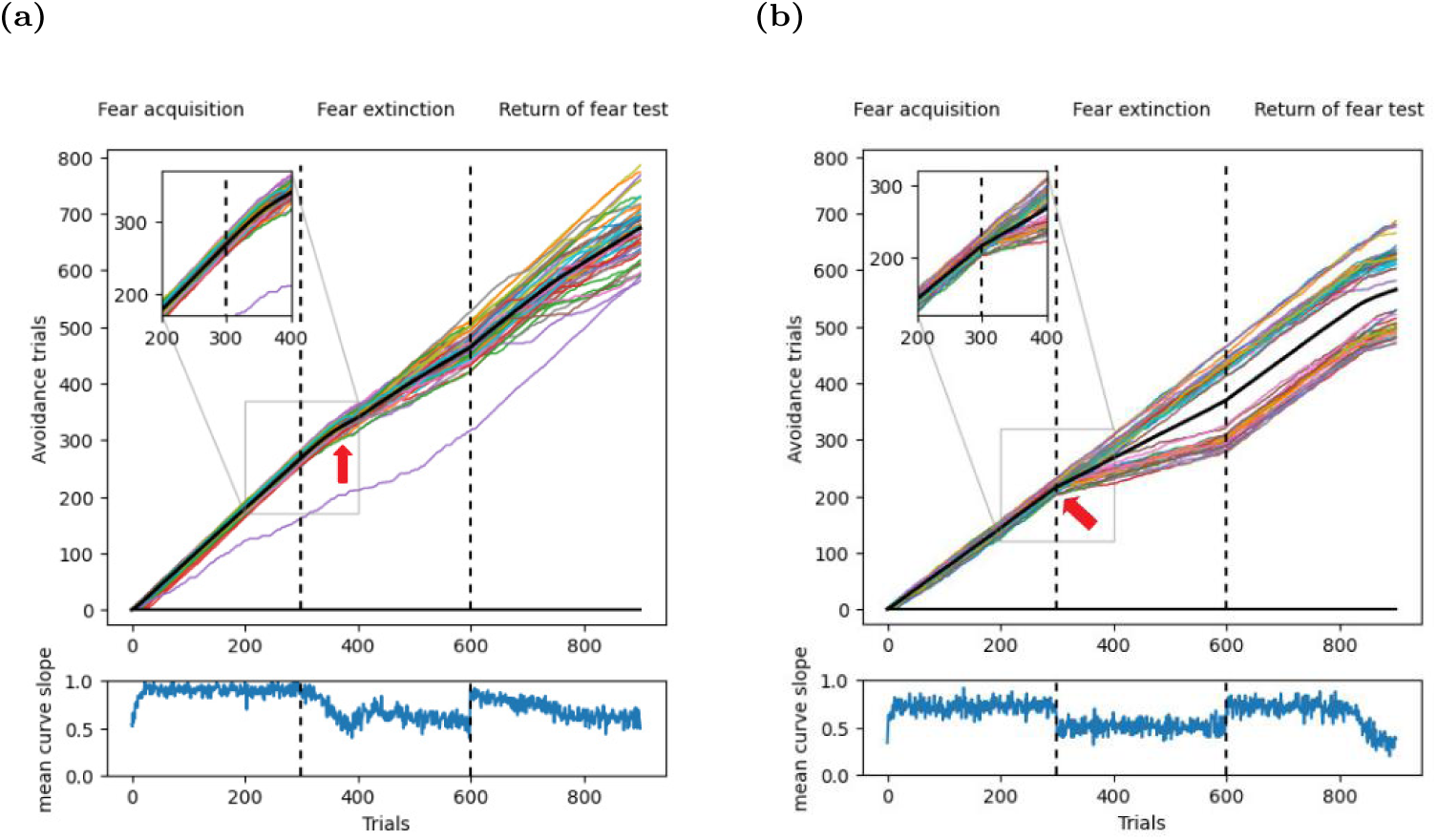
Dynamics of acquisition, extinction, and renewal-testing in T-maze. **(a)** Top: Cumulative response curves (CRCs) in an operant ABA renewal paradigm in the T-maze. Shown are 50 simulated CRCs (colored lines), and the mean CRC (black line). The inset shows the beginning of the extinction phase in context B, where the CRCs continue with the same slope as at the end of the acquisition phase in context A. This indicates that the association learned in context A generalized to context B. The red arrow points where the slopes start to change. Bottom: The slope of the mean CRC reveals rapid acquisition, gradual extinction learning, and renewal. **(b)** Top: In contrast to our model, Redish et al.’s model [17] changes its behavior immediately upon the switch from context A to context B (red arrow). Hence, this model does not generalize the learned association from context A to context B. In addition, one group of CRCs does not exhibit any extinction learning, and hence no renewal. Bottom: The slope of the mean CRC shows the sudden change in behavior at the onset of the extinction phase.

To test our model in an even more challenging task, we used an inhibitory avoidance learning paradigm in the shock zone maze, which combined complex spatial navigation, positive and negative rewards (see Methods), in an ABA renewal paradigm. In this task, agents had to navigate to a goal location in a 2-d environment along the shortest path possible, while at the same time avoiding a shock zone with an irregular shape

(Figs. 2e,2f). A trial was considered correct if the agent reached the goal without entering the shock zone on the way. This definition allowed us to use the CRC to quantify the trial-by-trial performance of simulated agents. The curve increased by one if a trial was correct and remained constant otherwise. That means, a slope of 1 in the CRC indicated perfect avoidance of the shock zone, and zero slope represented no avoidance at all.

Even in this complex learning task, the memory-intact agent in our model showed biologically realistic behavior. The mean CRC started with a slope of about 0.0 in the acquisition phase (Fig. 6a). After ≈ 100 trials, the CRC slope increased to about 0.8, as the agent learned to circumnavigate the shock zone. At the beginning of the extinction phase, the agent continued to avoid the shock zone, even though shocks were no longer applied and even though the context had changed from A to B. However, by the end of the extinction phase, the agent learned to reach the goal faster by traversing the shock zone, as indicated by a CRC slope of about 0.0. When the context was switched back to context A at the onset of the test phase, the CRC slope rose sharply, indicating renewal (Fig. 6a). Over the course of the test phase, the agents learned that the shock zone was no longer active in context A and that it could be safely traversed, causing a smooth drop of the CRC slope to about 0.0 in the final trials. These results showed that the agents in our model could learn context-dependent behavior even in a highly complex operant learning task in a two-dimensional environment.

**Fig 6.**
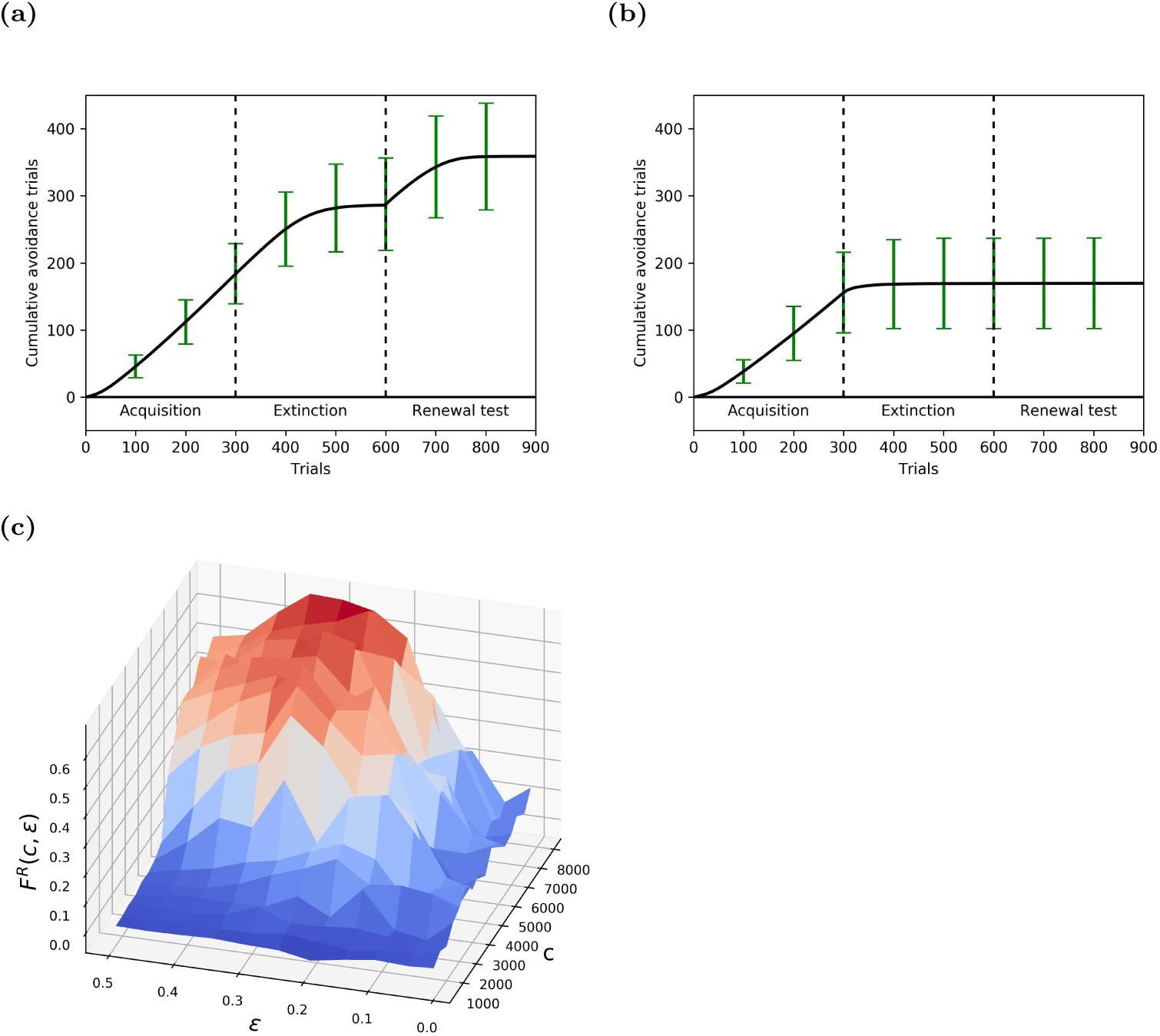
Learning dynamics of inhibitory avoidance in simulated ABA renewal. **(a)** Cumulative response curve (CRC) in the shock zone maze averaged across 1000 memory-intact agents (error bars indicate std). Agents showed clear evidence for extinction, i.e. flattening of CRC at the end of the extinction phase, and renewal, i.e. positive slope at the beginning of the renewal phase. **(b)** By contrast, memory-impaired agents showed faster extinction (flattening of CRC) and no renewal (flat CRC at onset of renewal testing). **(c)** Renewal strength *F*^*R*^ (*c, ϵ*) as a function of the capacity of the experience replay memory *c* and the *ϵ*-greedy exploration parameter. The renewal strength increases with *c*; note that the largest renewal effect is observed along the ridge for *c* > 4000 and 0.25 < *ϵ* < 0.35.

**Fig 7.**
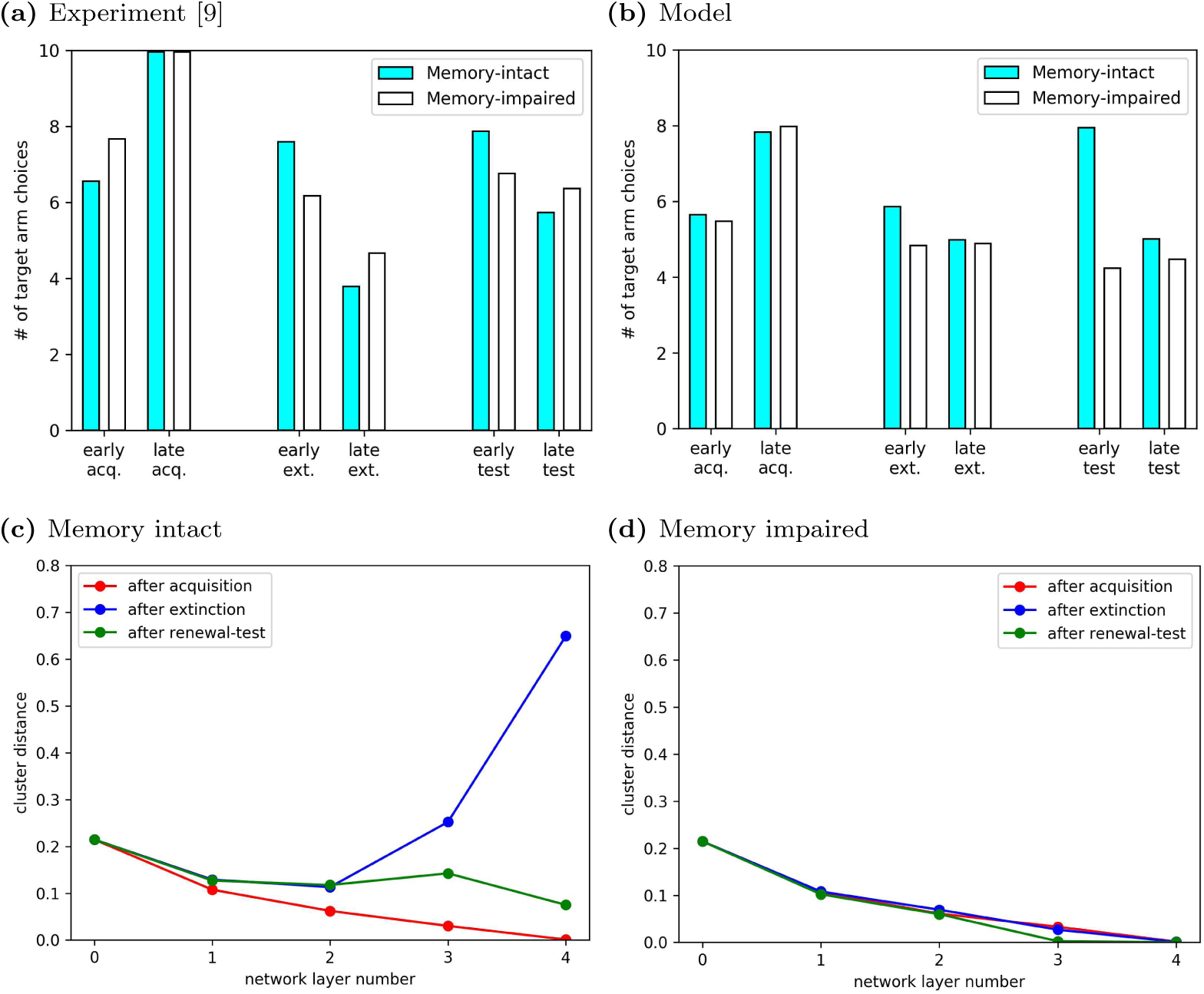
ABA renewal arises because experience replay drives the emergence of two distinct context representations. **(a)** André and Manahan-Vaughan (2016) showed that renewal was reduced by a pharmacological manipulation that interfered with synaptic plasticity in the hippocampus. (Adapted from Fig. 1 in [9].) Renewal score in experimental animals: *S*^*R*^ ≈ 0.45, renewal score in control animals: *S*^*R*^ ≈ 1.08. **(b)** In our model, agents with intact experience replay memory showed ABA renewal in the T-Maze task, i.e. an increase in response rate in early test despite the continued absence of reward (*S*^*R*^ ≈ 0.59). By contrast, memory-impaired agents did not exhibit renewal and had a lower renewal score (*S*^*R*^ ≈ −0.14). **(c)** Cluster distances between network representations of context A and B in the T-maze for memory-intact agents. Results were averaged over 1000 runs of our model. The representation of contexts A and B in higher layers of the deep neural network (*l* ≥ 3) formed distinct clusters that were well separated. They were learned in the extinction phase, since the distinction was not present at the end of the acquisition phase. **(d)** By contrast, memory-impaired agents did not show distinct context representations in any learning phase.

### The Role of Experience Replay in Learning Context-dependent Behavior

We next explored the role of experience replay in the emergence of context-dependent behavior in our model agent. To this end, we repeated the above simulation with a memory-impaired agent. The memory-impaired agent learned in much the same way as the memory-intact agent did during acquisition (Fig. 6b), but exhibited markedly different behavior thereafter. First, extinction learning was faster in the memory-impaired agent, consistent with experimental observations. This result might seem unexpected at first glance, but it can be readily explained. With a low memory capacity, experiences from the preceding acquisition phase were not available during extinction and the learning of new reward contingencies during extinction led to catastrophic forgetting in the DQN [24]. As a result, the previously acquired behavior was suppressed faster during extinction. Second, no renewal occurred in the test phase, i.e., the CRC slope remained about zero at the onset of the test phase. This is the result of catastrophic forgetting during extinction. This result is consistent with experimental observations that hippocampal animals do not exhibit renewal and suggests that the crucial function of the hippocampus in renewal is its role in memory replay. Our results suggest that ABA renewal critically depends on storing experiences from the acquisition phase in experience memory and making them available during extinction learning.

The agent’s learning behavior in our model crucially depended on two hyperparameters, namely the capacity of the experience replay memory *c*, and the exploration parameter *ϵ*. To analyze the impact of these hyperparameters, we performed and averaged 20 runs in the shock zone maze for different parameter combinations of *c* and *ϵ*. Let *S*^−^ (*c, ϵ*) and *S*^+^ (*c, ϵ*) be the slopes of the mean CRCs in the last 50 steps of extinction and first 50 steps of the test phase, respectively. In the ideal case of extinction followed by renewal: *S*^−^ (*c, ϵ*) = 0.0 and *S*^+^ (*c, ϵ*) = 1.0. We therefore defined the renewal strength for each combination of hyperparameters as

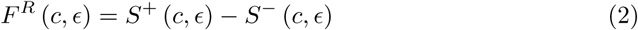

which is near 1 for ideal renewal and near 0 for no renewal. As expected from our results on memory-impaired agents above, the renewal strength was near zero for low memory capacity (Fig. 6c). It increased smoothly with increasing memory capacity *c*, so that a memory capacity *c* > 4000 allowed the agents to exhibit renewal reliably. The renewal strength displayed a stable ridge for *c* > 4000 and 0.25 < *ϵ* < 0.35. This not only justified setting *ϵ* = 0.3 in our other simulations, but also showed that our model is robust against small deviations of *ϵ* from that value.

The renewal strength was very low for small values of *ϵ* regardless of memory capacity. This points to an important role of exploration in renewal. With a low rate of exploration in the extinction phase, the model agent persevered and followed the path learned during acquisition. This led to *S*^−^ (*c, ϵ*) *S*^+^ ≈ (*c, ϵ*), and hence *F*^*R*^ (*c, ϵ*) ≈ 0, and explained why the exploration parameter has to take relatively large values of 0.25 < *ϵ* < 0.35 to generate robust renewal.

### Emergence of Distinct Neural Representations for Contexts

Having shown that our model learned context-dependent behavior in complex tasks, we turned our attention to the mechanism underlying this behavior in a specific experiment. We simulated an experiment in appetitive, operant extinction learning that found that increasing the dopamine receptor agonists in the hippocampus had a profound effect on ABA renewal in a simple T-maze [9] (Fig. 5a). The model agent received rewards for reaching the end of either side arm, but the reward was much larger in the left (target) arm. Like in the experimental study [9], we quantified the performance of the simulated agents by the number of trials, out of 10, that the agents chose the target arm at different points of the simulated experiment: *early acq.*,*late acq.*, *early ext.*, *late ext.*, *early test*, *late test*. In the acquisition phase in context A, memory-intact agents initially chose randomly between the left and right arm, but by the end of acquisition they preferentially chose the target arm (Fig. 5b, blue filled bars). In the extinction phase in context B, agents initially continued to prefer the target arm, i.e. they generalized the association formed in context A to context B, but this preference attenuated by the end of extinction. Critically, when the context was then switched back to the acquisition context A in the test phase, agents preferred the target arm again, even though it was no longer associated with a larger reward. This is the hallmark of ABA renewal. The renewal score *S*^*R*^ = (*early test/late ext.*) − 1.0 indicated the relative strength of the renewal effect in the T-maze experiments. *S*^*R*^ ≈ 0.59 for the memory-intact agent. Since there was no special reward for preferentially choosing the target arm, the agent stopped doing so by the end of the test phase.

The memory-impaired agent showed same acquisition learning as the memory-intact agent, but exhibited faster extinction learning and no renewal (*S*^*R*^ ≈ − 0.14, Fig. 5b, unfilled bars). These results are consistent with our modeling results in the inhibitory avoidance task discussed above and the experimental results from rats [9].

We hypothesized that context-dependent ABA renewal emerged in memory-intact agents because the higher layers of the DQN developed distinct representations of contexts A and B, which would allow the DQN to associate different reward contingencies with similar inputs. We therefore studied the evolution of context representations in the DQN based on the cluster distances between the representations of the two contexts (see Methods). Since the contexts A and B were distinguished by different illumination colors, the cluster distance in the input layer, *d*_0_, was nonzero (Fig. 5c). At the end of acquisition (red curve), the agents had been exposed only to visual inputs from context A, so there was no benefit for the DQN to represent the contexts A and B differently. Therefore, the difference in the context representations in the input layer was gradually washed out in higher layers, i.e., *d*_1_,…, *d*_4_ < *d*_0_. However, after extinction (blue curve), the replay memory contained observations from both contexts A and B, which were associated with different reward contingencies. This required that the higher layers of the DQN discriminate between the contexts, and *d*_3_, *d*_4_ > *d*_0_. The network’s ability to represent the two contexts differently, and thus maintain different associations during the extinction phase, gave rise to renewal in the subsequent testing phase (green curve), where the experience replay memory encoded novel observations from context A, yet still contained residual experiences from context B. Reward contingencies were identical in both contexts, and there was no longer a benefit to maintaining distinct context representations, hence, *d*_3_, *d*_4_ ≲ *d*_0_. Note that *d*_3_ and *d*_4_ were also much smaller at the end of the test phase than they were at the end of extinction, and would eventually vanish if the renewal test phase was extended.

As expected, cluster distances for the DQN in memory-impaired agents (Fig. 5d) did not change between the phases of the experiment and remained at low values throughout. This was the consequence of the reduced experience replay memory capacity and accounted for the lack of renewal in memory-impaired agents. Since the agents had no memory of the experiences during acquisition in context A, the DQN suffered from catastrophic forgetting [24] when exposed to a different reward contingency in context B and therefore the DQN could not develop separate representations of the two contexts during extinction.

## Discussion

We have developed a computational model to study the emergence of context-dependent behavior in complex learning tasks within biologically realistic environments based on raw sensory inputs. Learning is modeled as deep Q-learning with experience replay [15]. Here, we apply our model to a sparsely researched topic: modeling of operant extinction learning [3]. We found that our model accounts for many features that were observed in experiments, in particular, in context-dependent ABA renewal, and the importance of the hippocampus in this processes. We assumed in our model that the primary function of the hippocampus is to store experiences and replay them later. Our model has several advantages relative to previous models. Compared to Redish et al.’s model of operant extinction learning [17], our model learned faster on the linear track, and showed more plausible extinction learning in that our model generalized from the acquisition to the extinction context. Further, our model allowed for the investigation of internal representations of the inputs and, hence, the comparison to neural representations. Finally, our model makes two testable predictions: Context-dependent behavior arises due to the learning of clustered-representations of different contexts, and context representations are learned based on memory replay driven by the hippocampus. We discuss these predictions in further detail below.

### Learning context representations

One common suggestion in the study of learning is that stimuli can be distinguished *a priori* into discrete cues and contextual stimuli based on some inherent properties [25]. For instance, cues are discrete and rapidly changing, whereas the context is diffuse and slowly changing. Prior models of extinction learning, e.g. [16, 17], rely on external signals that isolate sensory stimuli and the current context. They maintain different associations for different contexts and switch between associations based on the explicit context signal. Since this explicit context switching is biologically unrealistic, Gershman et al. suggested a latent cause model in which contexts act as latent causes that predict different reward contingencies [26]. When expected rewards do not occur, or unexpected rewards are delivered, the model infers that the latent cause has changed. However, the model still depends on explicit stimulus signals, and requires complex Bayesian inference techniques to create and maintain the latent causes.

An alternative view is that there is no categorical difference between discrete cues and contextual information during associative learning, i.e., all sensory inputs drive learning to some extent [14]. Two lines of experimental evidence support this learning hypothesis. First, the learning of context can be modulated by attention and the informational value of contexts [27]. This finding suggests that contextual information is not fixed and what counts as contextual information depends on task demands. Second, the context is associated with the US to some degree [28], instead of merely gating the association between CS and US. There are suggestions that the association strength of the context is higher in the early stage of conditioning than later, when it becomes more apparent that the context does not predict the US [28]. However, when context is predictive of CS-US pairing, the CR remains strongly context-dependent after multiple learning trials [29].

In line with the learning view, the internal deep Q-network in our model received only raw camera inputs and no explicit stimuli or context signaling. The model autonomously adapted the network weights to given learning tasks based on reward information collected during exploration and from the experience replay memory. By analyzing the model’s internal deep Q-network, we found that distinct context representations emerged spontaneously in the network activations over the course of learning, if the agent was exposed to two contexts with different reward contingencies.

### The neural representation of context

The hippocampus has been suggested to play a role in the encoding, memorization, and signaling of contexts, especially in extinction learning [3, 11]. However, there is no conclusive evidence to rule out that the hippocampus is involved in the renewal phenomenon primarily due to its other well-established role in general-purpose memory storage and retrieval [30–32]. In our model, the emergence of distinct context representations critically depended on experience replay, suggesting that experience replay facilitates the formation of stable memory representations – as predicted by neuroscientific studies [19, 33]. We propose that episodic memory incidentally stores observations and actions in an environment for later use. According to this view, the spatial representations so often observed and studied in the hippocampus [25, 34] would then be a result of the spatial information that is present in the inputs to the hippocampus [35], and would not constitute evidence for context memorization in the hippocampus, as proposed earlier [25].

In line with previous suggestions [6, 10], we therefore propose that contexts are represented, if at all, in brain regions other than the hippocampus and that learning of these representations is facilitated by memory of experiences (inputs, actions and rewards), which are stored with the help of the hippocampus. To test this hypothesis, our model, unlike previous computational models of extinction learning, cleanly delineates the simulated hippocampal storage system (experience replay memory) from other cortical regions (deep Q-network). As the network learns abstract representations of the visual inputs, the lower layers most likely correspond to visual areas in the cortex, while the higher layers serve functions similar to those served by the prefrontal cortex (PFC).

We propose that this hypothesis could be tested using high-resolution calcium imaging [36] of the PFC in behaving rodents. Our model predicts that different contexts are encoded by PFC population activity that form well-separated clusters by the end of extinction learning, allowing the animals to disambiguate contexts, and leading to renewal. In hippocampal animals, however, we expect that novel contexts are not encoded in separate PFC clusters, but instead their representations in the PFC population will overlap with those of previously acquired contexts. Furthermore, our model suggests that context representations in the brain are highly dynamic during renewal experiments, adapting to the requirements of each experimental phase. Distinct context representations would therefore arise in control animals over the course of extinction learning, and not necessarily be present from the start.

### The role of the hippocampus in renewal

Our hypotheses about contextual representations discussed above naturally beg the question: What role does the hippocampus play in renewal? The experimental evidence leaves no doubt that the hippocampus plays a critical role in renewal [3, 7, 11, 37]. Based on our modeling results, we propose that the hippocampus is involved in renewal because it stores and replays experiences. In our model, replay of experiences from the acquisition phase was critical to learn distinct context representations in the extinction phase. This is consistent with observations in humans [38]. Only subjects whose activation in the bilateral hippocampus was higher during extinction in a novel context than in the acquisition context showed renewal later. With limited experience memory, learning during extinction in our model overwrites previously acquired representations, also known as catastrophic forgetting. This in turn prevented renewal from occurring when the agent was returned to the acquisition context. Note that the experience memory in our model by itself does not extract any information about context from the inputs. To summarize, our model predicts that impairing the hippocampus pre-extinction weakens renewal, while impairing the hippocampus post-extinction should have no significant influence on the expression of renewal.

Both of these predictions are supported by some experimental studies, and challenged by others. Regarding the first prediction, some studies have confirmed that hippocampal manipulations pre-extinction have a negative impact on ABA renewal. For instance, ABA renewal was disrupted by pharmacological activation of dopamine receptors prior to extinction learning [9]. By contrast, other studies suggest that disabling the hippocampus pre-extinction has no impact on later renewal. For instance, inactivation of the dorsal hippocampus with muscimol, an agonist of GABA_A_ receptors [39], or scopolamine, a muscarinic acetylcholine receptor antagonist [8], prior to extinction learning did not weaken ABA renewal during testing. The apparent contradiction could be resolved, if the experimental manipulations in [8, 39] did not interfere with memories encoded during acquisition, as other manipulations of the hippocampus would, but instead interfered with the encoding of extinction learning. As a result, the acquired behavior would persist during the subsequent renewal-test. Indeed, activation of GABA_A_ receptors, or the inactivation of muscarinic receptors reduce excitation of principle cells, thereby changing signal-to noise ratios in the hippocampus. We hypothesize that, as a result, animals exhibited reduced extinction learning following scopolamine [8] or muscimol [39], consistent with impaired encoding of extinction. By contrast, dopamine receptor activation during extinction learning [9], did not reduce extinction learning, but impaired renewal. The results of the latter study are more consistent with the results of our model when memory capacity is reduced pre-extinction: extinction learning was slightly faster and renewal was absent.

In support of our second prediction, a study showed that antagonism of β-adrenergic receptors, which are essential for long-term hippocampus-dependent memory and synaptic plasticity [40], post-extinction had no effect on ABA renewal [41]. By contrast, several studies suggested that lesioning the hippocampus post-extinction reduces renewal of context-dependent aversive experience. For instance, ABA renewal in the test phase was abolished by excitotoxic [37] and electrolytic [7] lesions of the dorsal hippocampus after the extinction phase. More targeted lesions of dorsal CA1, but not CA3, post-extinction impaired AAB renewal [6]. The authors of these studies generally interpreted their results to indicate that the hippocampus, or more specifically CA1, are involved in retrieving the contextual information necessary to account for renewal. However, damage to these subregions can be expected to have generalized effects on hippocampal information processing of any kind. Another possibility is that post-extinction manipulations, which were made within a day of extinction learning, interfered with systems consolidation. This is very plausible in terms of dopamine receptor activation [9] as this can be expected to promote the consolidation of the extinction learning event. To model this account in our framework, one would have to split the learning and replay during extinction phase, which occur in parallel in our current implementation, into two separate stages. Only new learning would occur during the extinction phase, while experience replay (of the acquisition phase) would occur during sleep or wakeful quiescence after the extinction phase, as suggested by the two-stage model of systems consolidation. In this case, the associations formed during acquisition would be initially overwritten during extinction learning, as they were in our simulations without experience memory, and then relearned during the replay/consolidation period [19, 20]. If systems consolidation was altered during this period, extinction learning would still be retained, but the association learned during acquisition would be overwritten and hence there would be no renewal. In line with this interpretation, recent findings suggest that extinction learning and renewal engage similar hippocampal subregions and neuronal populations [13], thus, suggesting that manipulating the hippocampus can have an impact on prior learned experience and thus, subsequent renewal.

In conclusion, we have shown that, in an ABA renewal paradigm, contextual representations can emerge in a neural network spontaneously driven only by raw visual inputs and reward contingencies. There is no need for external signals that inform the agent about cues and contexts. Since contextual representations might be learned, they might also be much more dynamic than previously thought. Memory is critical in facilitating the emergence of distinct context representations during learning, even if the memory itself does not code for context representation per se.

## Acknowledgments

Funded by the Deutsche Forschungsgemeinschaft (DFG, German Research Foundation) – project number 316803389 – through SFB 1280, projects A04 (D.M.V.), A14 (S.C.) and F01 (S.C.).

## Footnotes

1 Chollet, F. et al., 2015, https://keras.io

2 Plappert, M., 2016, https://github.com/keras-rl/keras-rl

## Notes

### Competing Interest Statement

The authors have declared no competing interest.

